# Contribution of Mesenchymal-like and Epithelial Cellular Subsets to Chemotherapy Resistance in Triple-Negative Breast Cancer

**DOI:** 10.1101/2025.11.01.685128

**Authors:** Ngoc B. Vuong, Olga Y. Korolkova, Michael Izban, Nobelle I. Sakwe, Antonisha R. McIntosh, Destiny Ball, Perrin Black, Alayjha Edwards, Billy R. Ballrad, Samuel E. Adunyah, Amos M. Sakwe

**Author notes:** **Corresponding Author:** Amos M Sakwe, Department of Biomedical Sciences, Meharry Medical College, Nashville, Tennessee, 37208, USA.

## Abstract

**Background/Objectives:** Triple-negative breast cancer (TNBC) tumors are typically heterogeneous, predominantly epithelial tissues with discrete patches of mesenchymal-like TNBC cells, that differ in their invasiveness, proliferation potential and response to treatment. However, the contribution of mesenchymal-like and epithelial TNBC cells in the persistence of chemotherapy resistant disease remains poorly understood.

**Methods:** Mesenchymal-like and epithelial TNBC cell types were detected by multiplex fluorescent immunohistochemistry using antibodies against vimentin, Ki67, and Annexin A6 (AnxA6). Chemotherapy drug resistant mesenchymal-like and epithelial TNBC cell populations were established by pulse exposure and stepwise dose escalation and validated by 3D cultures and unbiased antibody arrays.

**Results:** Analysis of the response of stage IV TNBC tumors treated with six common chemotherapy regimens resulted in 36% complete response and 64% partial response with residual tumor sizes ranging from 0.5 to 37.0 mm. Treatment of TNBC cells with chemotherapy agents led to distinct resistance signatures including downregulation of survivin and upregulation of M-CSF and CXCL8/IL-8 in model mesenchymal-like, and upregulation of CCL2/MCP-1, CTSS and DKK-1 in model epithelial TNBC cells. The inhibitory phosphorylation of GSK-3β (p-S9) increased in paclitaxel resistant epithelial cells but decreased in resistant mesenchymal-like TNBC cells. Finally, chemotherapy resistance also activated p90 ribosomal S6 kinases (RSK1/2) in both cell types, while activation of mitogen- and stress-activated kinases (MSK1/2) was only observed in chemotherapy resistant epithelial TNBC cells.

**Conclusions:** These data reveal that chemotherapy resistance of epithelial and mesenchymal-like TNBC cellular subsets upregulated distinct profiles of proinflammatory and immune cell chemotactic cytokines and modulated the activities of GSK-3β, p90 RSK1/2 and the related MSK1/2. Targeting these factors and/or the associated signaling pathways may help overcome chemotherapy resistance in TNBC.

**Graphical Abstract:** 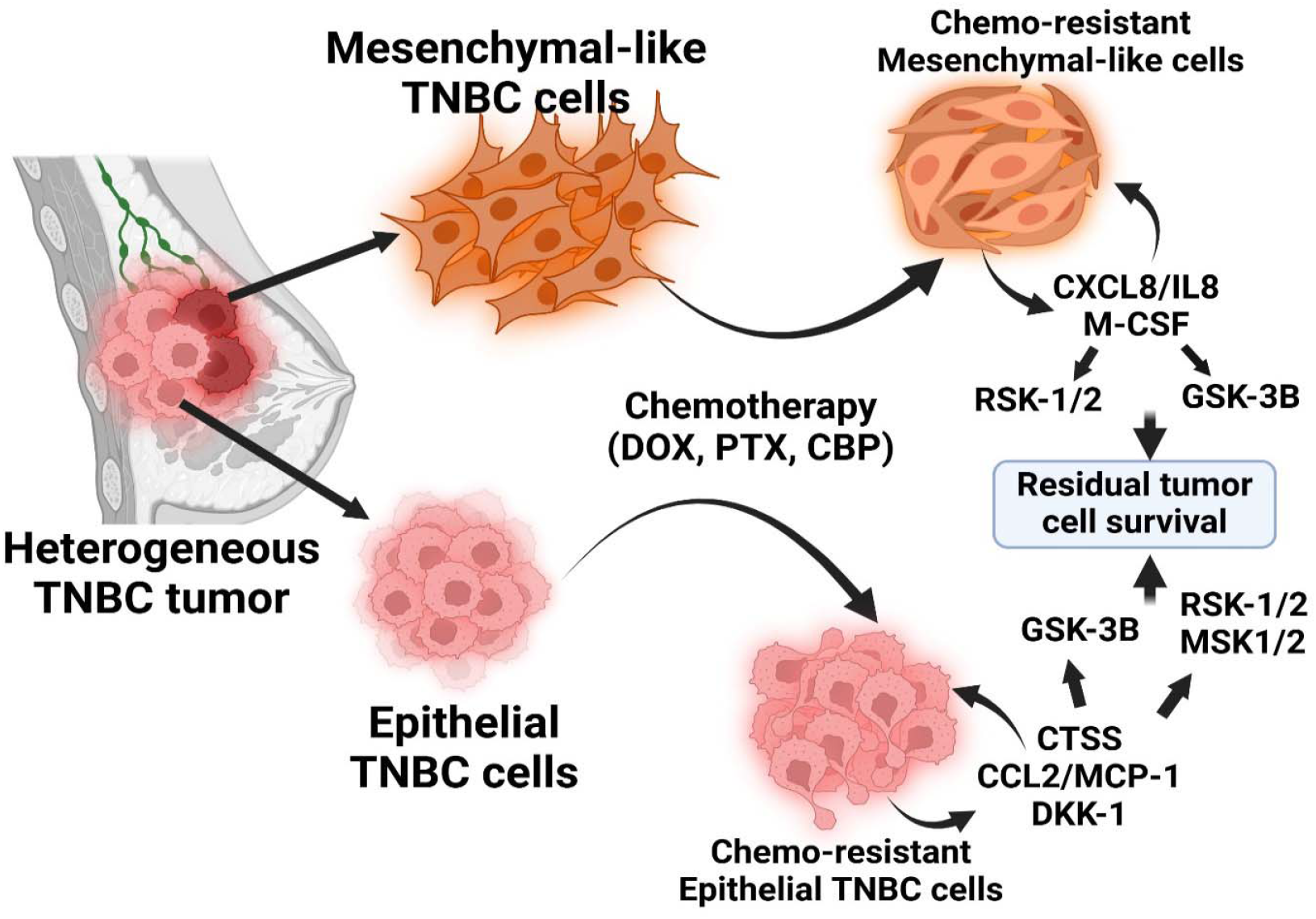

TNBC tumors are typically heterogeneous, predominantly epithelial tissues with discrete patches of mesenchymal-like TNBC cells, that differ in their invasiveness, proliferation potential and response to treatment. This study reveals that Chemotherapy resistance in epithelial and mesenchymal-like TNBC cells is maintained by distinct signatures of proinflammatory and immune cell chemotactic cytokines, and activities of GSK-3β and p90 ribosomal S6 kinases.

## Introduction

Triple negative breast cancer (TNBC) represents 15-20% of incident breast cancers, but accounts for nearly 40% mortality within the first 5 years after diagnosis, compared to other breast cancer subtypes (1). This highly aggressive breast cancer subtype is characterized by the absence of estrogen and progesterone receptors, and HER2 amplification. The lack of these receptor makes TNBC tumors particularly challenging to treat, as they do not respond to either hormonal therapies or HER2-targeted treatments (2). Consequently, chemotherapy remains the treatment of choice for TNBC, and despite initial responsiveness, TNBC rapidly relapses and becomes resistant to subsequent treatments (3-5).

Many studies have used gene expression profiling to demonstrate that the diverse response of TNBC tumors to treatment is partly due to tumor heterogeneity. This includes distinct TNBC molecular subtypes which now comprise mesenchymal-like (MSL), basal-like immune activated (BL1/BLIA), basal-like immune suppressed (BL2/BLIS), and luminal androgen receptor positive (LAR) (6-9). These TNBC molecular subtypes are also known to exhibit overlapping alterations in DNA repair pathways, metabolic pathways, immune checkpoint components as well as significant differences in the composition of the tumor microenvironment (10). Although these classifications provide insight into personalized treatment options, the overlapping alterations invariably contribute to the varied responses of TNBC tumors to chemotherapy and subsequent development of resistance (11). Therefore, the impact of these classifications on clinical outcomes remains unpredictable.

The complexity of TNBC tumors like most breast cancers also includes multiple breast cancer subtypes (8, 12) and morphologically distinct mesenchymal-like and epithelial cell types that exhibit distinct biological behaviors including invasiveness and therapeutic susceptibilities (13-15). The impact of intra-tumoral heterogeneity on treatment resistance is not only defined by evolution of distinct phenotypic cell populations as the disease progresses (16), but also how the cellular subsets with distinct intrinsic drug sensitivities co-exist within the same tumor (17). Epithelial TNBC cells typically retain cell-cell adhesion properties and lead to larger but less aggressive tumors. Mesenchymal-like TNBC cells on the other hand, are characterized by their enhanced migratory and invasive properties, often associated with epithelial-mesenchymal transition (EMT), stem cell-like properties and resistance to apoptosis (18, 19). Given that typical TNBC tumors consist of distinct proportions of epithelial cells and tumor cells at various stages of EMT, a better understanding of subtle molecular differences between these phenotypically distinct TNBC cell types following chemotherapy is essential to eventually develop novel and more effective treatment regimens.

The impact of phenotypic diversity measured by Simpson’s score does not always correlate with genetic intra-tumor heterogeneity measured by the MATH index especially on responses to treatment (20). Classification of TNBC into proliferative basal-like (here in denoted as epithelial) and the more invasive mesenchymal-like subsets relies on cell morphology and expression of phenotypic molecular markers such as the epithelial E-cadherin and the mesenchymal vimentin (2, 18, 21). Both markers are known to be heterogeneously expressed and change with cancer progression, requiring the use of other cell proliferation markers such as Ki67, or other markers of invasiveness. Detection of Ki67 and Annexin A6 (AnxA6), a multifunctional scaffolding protein that is critical in cell proliferation, survival, migration, membrane repair and drug resistance (22-25), has previously been reported to delineate AnxA6-^high^ invasive from AnxA6-^low^ rapidly growing TNBC cell lines and patient derived xenograft models (26). While this supports the detection of AnxA6 in TNBC tumors as a marker of invasiveness, its association with the expression of vimentin in TNBC cells remains unclear.

The goal of this study was to determine the contribution of mesenchymal-like and epithelial TNBC cellular subsets to chemotherapy resistance. We assessed the response of TNBC patients to six different chemotherapy regimens, established the detection of epithelial and mesenchymal-like cells within the tumors, and identified molecular differences between residual chemotherapy resistant phenotypically distinct mesenchymal-like and epithelial TNBC cell types.

## Materials and Methods

### Cell culture

The mesenchymal-like BT-549 cells (ATCC, HTB-122) and epithelial-like MDA-MB-468 (ATCC, HTB-132) cells were obtained from the American Type Culture Collection (Manassas, VA). Only the early (< 5) passages were used in experiments. BT-549 cells were cultured in DMEM/F12 (Thermo Fisher Scientific, Waltham, MA) medium supplemented with 10% fetal bovine serum (FBS), 5% sodium bicarbonate and 1% Penicillin/Streptomycin. MDA-MB-468 cells were cultured in Leibovitz’s L-15 medium (Thermo Fisher Scientific, Waltham, MA) supplemented with 10% fetal bovine serum (FBS), 5% sodium bicarbonate and 1% Penicillin/Streptomycin. Cells were maintained at 37 °C with 5% CO2 in a humidified incubator. Media were changed every 2-3 days and cells were passaged by using TrypLE cell dissociation reagent (Gibco), and the cells were regularly checked for mycoplasma contamination using mycoplasma detection kit from InvoivoGen US (San Diego, CA).

### Establishment of resistant TNBC cell lines

Doxorubicin HCl (DOX) (NSC 123127), Paclitaxel (PTX) (NSC 125973), and Carboplatin (CBP) (NSC 241240) were procured from Selleck Chemicals LLC (Houston, TX) and used to establish drug-resistant BT-549 and MDA-MB-468 cells using the pulse exposure and stepwise dose escalation method as previously described for cisplatin resistant ovarian cancer cells (27). Briefly, the cells were treated with each chemotherapy agent by continuous exposure of gradually increasing concentrations of the drugs over a period of 6 months. BT-549 or MDA-MB-468 cells were cultured in the drug-containing media for 48 h, allowed to recover in drug-free media until 80% confluency, and the treatment cycle repeated with the same concentration of drug. A new cycle began with maintaining the cells in media containing a slightly higher concentration of the drugs until concentrations at which the cells could not survive due to cytotoxicity.

### Cell proliferation

Cells were seeded at a density of 5000 cells per well in an 8-well chamber Millicell EZ slide (Sigma-Aldrich, St. Louis, MO) for 48 h. Cells were stained using the baseclick GmbH EdU HTS assay kit as described by the manufacturer (Sigma-Aldrich, St. Louis, MO). Briefly, cells were incubated with EdU for 3 h, then fixed and permeabilized in cold methanol at −20 °C followed by EdU detection using the red fluorescent Cyanine 5 (Cy5) dye and cell imaging (546/579nm Ex/Em).

### Cell viability and clonogenic assays

For cell viability assays, cells were seeded in 96-well plates at a density of 5000 cells per well. Cells were treated with the indicated drugs for 72 h and then incubated with PrestoBlue™ Cell Viability Reagent (Invitrogen, CA) for 1 h at 37 °C in a humidified incubator. The viability of the cells was measured by fluorescence intensity (530/590 Ex/Em) by using a Biotek Synergy HT Microplate Reader (Hampton, NH). For clonogenic assays, cells were plates in 24 well ultralow attachment plates in complete medium supplemented with 2% Matrigel and the indicated drugs. Cells were cultured for up to 10 days and brightfield images were captured using Accu-Scope EXI-410 inverted phase contrast microscope (Commack, NY) at 10x magnification.

### Immunofluorescence

Cells were seeded at a density of 5000 cells per well in the 8-well chamber Millicell EZ slide (Sigma-Aldrich, St. Louis, MO) for 48 h and immunofluorescence was carried out as previously described (28). Cells were fixed with cold methanol at room temperature (RT) for 20 min, followed by blocking with 1% bovine serum albumin (BSA) in phosphate buffered saline (PBS) for 1 h. Cells were then incubated for 1 h with the following primary antibodies Annexin VI (Santa Cruz Biotechnology, Dallas, TX), Vimentin (Santa Cruz Biotechnology, Dallas, TX) and Ki67/MKI67 (R&D Systems, Inc., Minneapolis, MN). Cells were then stained using Alexa Fluor™ Plus 488 conjugated Donkey anti-Rabbit IgG (Invitrogen, USA) and Alexa Fluor™ 568 conjugated Donkey anti-Mouse IgG (Invitrogen, USA) for 45 min and counterstained with ProLong™ Gold Antifade mounting fluid with DAPI (Invitrogen) for 10 min, at RT. Imaging was performed using BZ-X710 Imaging Reader (Keyence, Itasca, IL).

### Immunocytochemistry and multiplex fluorescent immunohistochemistry

The use of clinical and pathological tissues from 22 stage IV TNBC patients in this study was approved by the Meharry Medical College Institutional Review Board (IRB) as exempt research. Slides containing formalin fixed paraffin embedded deidentified biopsies (primary treatment naïve) or residual chemotherapy resistant tumor tissues were obtained from the Vanderbilt Translational Pathology Shared Resource as previously described (26). The slides were processed for standard immunohistochemistry (IHC) and images were captured by using C2+ confocal microscope and NIS-Elements software (Nikon Europe B.V., The Netherlands). Multiplex fluorescent IHC was performed by using the tyramide signal amplification procedure as previously described (29).

Deparaffinized tissues were incubated in citrate buffer for antigen retrieval, then permeabilized and blocked with 3% hydrogen peroxide solution at 37°C for 20 min. Non-specific blocking with normal goat serum (Santa Cruz Biotechnology) was performed, followed by sequential incubation with the following primary antibodies: Annexin VI (Santa Cruz Biotechnology, Dallas, TX), Vimentin (Santa Cruz Biotechnology, Dallas, TX) and Ki67/MKI67 (R&D Systems, Inc., Minneapolis, MN). Incubation with each primary and the corresponding secondary antibodies was followed by the antigen retrieval step to remove unbound antibodies. After secondary antibody incubation, tissue slides were washed and then incubated with a tyramide-conjugated fluorescent dye (Ty-fluor), rinsed and mounted with coverslip in media containing DAPI for nuclear counterstaining. Whole slide imaging and analysis was performed at the Digital Histology Shared Resource at Vanderbilt University Medical Center. The fluorescent, immunostained tissue slides were imaged on an Aperio Versa 200 automated slide scanner (Leica Biosystems) at 20X magnification to a resolution of 0.323 µm/pixel. Tumor areas were mapped using Ariol Review software and the staining intensity of the images was performed using the Ariol software capable of triple and quadruple stained immunohistochemical (IHC) quantification.

### Proteome Profiler Antibody Arrays

The levels of 84 cancer-related proteins in cell lysates prepared from control and chemotherapy agent resistant cells were assayed by using the Proteome Profiler Human XL Oncology Array kit (R&D Systems; cat. no. ARY026). The activation status of 37 kinases was also assessed using the Proteome Profiler Human Phospho-Kinase Array Kit (R&D Systems; cat. no. ARY003C). Nitrocellulose membranes spotted in duplicate with capture antibodies against cancer-related proteins were incubated with cell lysates prepared from control and chemotherapy drug resistant cells and processed as recommended by the manufacturer (R&D Systems, Minneapolis, MN). The blots were developed using enhanced chemiluminescent reagent (Perkin Elmer), and the images were acquired with the ChemiDoc™Touch imaging system (Bio-Rad, Hercules, CA). Protein expression was determined by densitometry of the dot blots using protein array analyzer for ImageJ software and corrected with the positive controls in each array membrane, as recommended by the manufacturer.

### Western Blotting

Cells were cultured until ∼80% confluency, then scraped in ice-cold Hanks’ balanced salt solution (HBSS) containing 1 mM Ca^2+^ and 1 mM Mg^2+^. Cells pellets were resuspended in radioimmunoprecipitation assay (RIPA) lysis buffer (50 mM Tris-HCl, pH 7.4, 1% NP-40, 0.1% sodium deoxycholate, 150 mM NaCl, 1 mM EDTA) containing freshly added protease and phosphatase inhibitors, incubated on ice for 30 mins and centrifuged at >10000 xg for 10 min at 4 °C. Western blotting was carried out as previously described (30, 31), using primary antibodies against Annexin VI (AnxA6), Vimentin and Ep-CAM (Santa Cruz Biotechnology, Dallas, TX), total and phosphorylated RSK1/2 and MSK1/2 () and β-actin (Santa Cruz Biotechnology, Dallas, TX) as the loading control. The blots were revealed with enhanced chemiluminescent reagent (Perkin Elmer), imaged with the ChemiDoc™Touch gel imaging system (Bio-Rad, CA) and quantified by densitometry using ImageJ software.

### Statistical Analysis

Statistical analysis was performed using GraphPad Prism (San Diego, CA). For all other analyses, paired t-tests were used and a p-value < 0.05 was considered statistically significant.

## Results

### AnxA6 and Ki67 expression defined mesenchymal-like and epithelial phenotypes in TNBC tumors

We assessed the clinical response of stage IV TNBC patients previously described in Korolkova et al., 2020 (26) to six typical neoadjuvant chemotherapy combinations of Doxorubicin (Adriamycin), cyclophosphamide (Cytoxan), platinum-based cytotoxic compounds (cisplatin, carboplatin), abraxane (nab-paclitaxel), paclitaxel (Taxol), capecitabine (Xeloda). Out of 22 patients, 8 (36%) displayed complete response with no residual tumors (NRT), and 14 (64%) displayed progressive or stable disease or had residual tumors (RT) with tumor sizes varying from 0.5 to 35 mm (Figure 1A-C). Out of the six regimens, dense dose anthracycline/Cytoxan (AC) followed by taxol (ddAC,T) was the most effective at reducing

**Figure 1.**
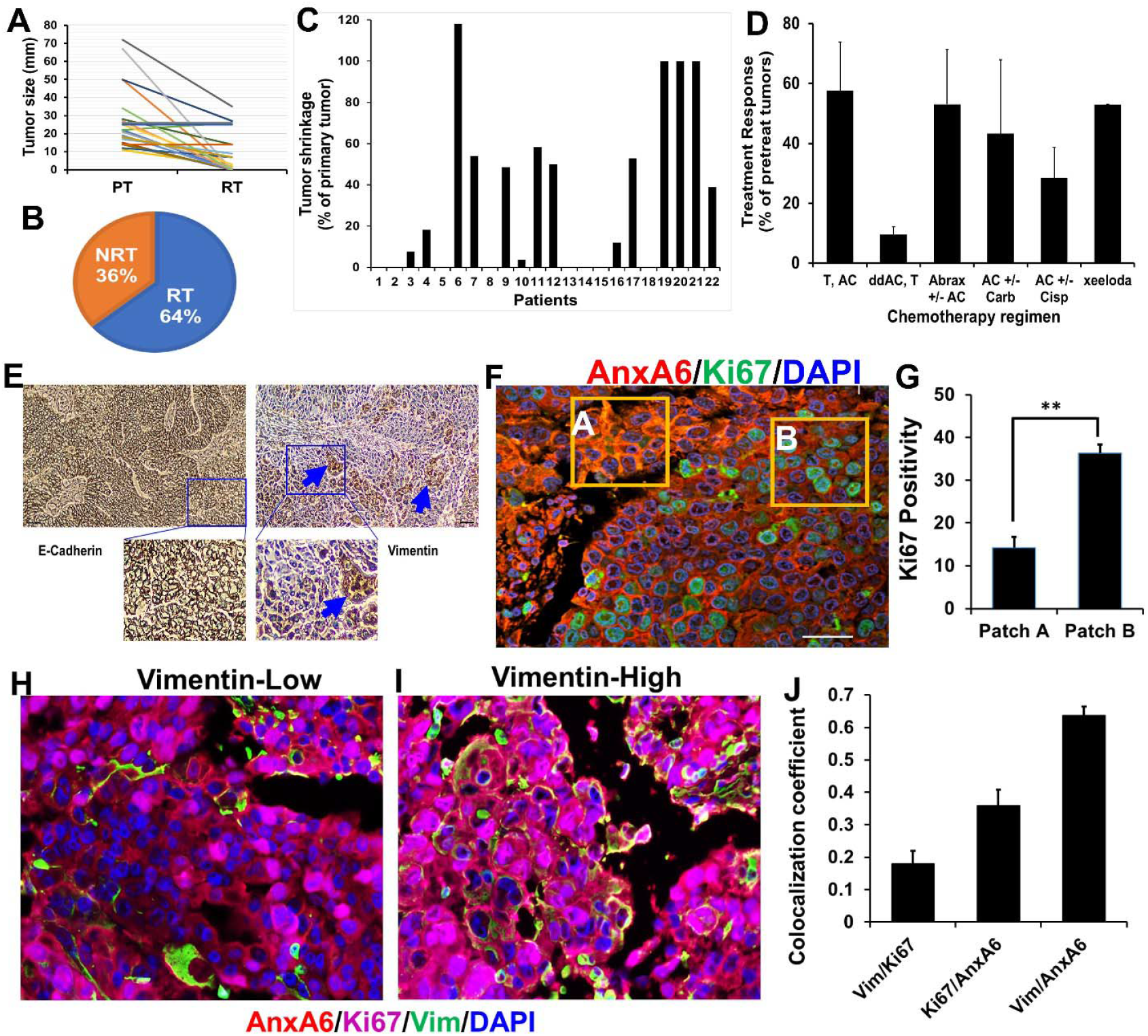
Response of stage IV TNBC tumors to different standard of care chemotherapy regimens and detection of epithelial and mesenchymal-like tumor cells. A-D) Analysis of the response and tumor sizes in TNBC patients be-fore and after six different neoadjuvant chemotherapy regimens. NRT = No Residual Tumor; RT = Residual Tumor; T = Taxol; AC = Adriamycin and Cytoxan; Cisp = Cisplatin; Carb = Carboplatin; dd AC = dose-dense Adriamycin; Abrax = Abraxane; Xeloda = Capecitabine. E) Detection of E-cadherin and vimentin in a typical TNBC tumor by immunohistochemistry (IHC). F-G) Validation of the inverse expression of AnxA6 (red), and cell proliferation marker Ki67 (green) in TNBC patient tissues and semi-quantification of the fluorescent intensity of Ki67 positivity. H-J) Relationship between AnxA6, Ki67 and vimentin in a typical TNBC tumor by multiplex fluorescent IHC. tumor size. Combinations of AC and T were the least effective, with larger recurrent tumors (Figure 1D).

To gain a better understanding of the diverse response to these treatment regimens, we assessed expression of E-cadherin and vimentin by IHC. This showed that a typical TNBC tumor is a predominantly epithelial tissue with discrete patches of mesenchymal-like TNBC cells (Figure 1E). This phenotype was further confirmed by double fluorescent IHC of the proliferation marker Ki67 and AnxA6, a calcium dependent membrane binding protein with high expression in invasive TNBC cells and low expression in proliferative TNBC cells (26). There were discrete tumor patches e.g. tumor area A with high expression of AnxA6 and low expression of Ki67 (AnxA6^hi^/Ki67^lo^) with ∼14% Ki67 positivity, while the bulk of the tumor represented by tumor area B showed AnxA6^lo^/Ki67^hi^ with >36%Ki67 positivity (Figure 1F-G). As shown in Figure 1H and I, vimentin staining was dispersed in vimentin-low patches and intensely expressed in more cells in vimentin high areas. In a typical TNBC tumor, the probability for co-expression of AnxA6 and vimentin is higher than with Ki67 or between AnxA6 and Ki67 (Figure 1J) and supports the use of vimentin and AnxA6 as bona fide markers of invasiveness. These data suggest that detection of Ki67, vimentin and AnxA6 can be used to assess phenotypic tumor heterogeneity and confirms that typical TNBC tumors are predominantly epithelial tissues with discrete areas of mesenchymal-like tumor cells.

We next assessed whether expression of AnxA6 and Ki67 is associated with vimentin expression in pre-treatment biopsies and chemotherapy residual disease tumors by multiplex immunofluorescence (Figure 2A) and the expression levels of these proteins in these tissues is shown as a heatmap (Fig. 2B). Analyses of the relationships between these proteins confirmed the reciprocal expression of AnxA6 and Ki67 in all tissues tested as previously reported (26). While co-expression of vimentin and AnxA6 was observed in some but not all tissues, expression of vimentin in both primary and chemotherapy residual strongly correlated with Ki67 (r = 0.9270, p = 0.0001), but not with AnxA6 (r = 0.1967, p = 0.4652). Despite the reciprocal expression of AnxA6 and Ki67, the expression of AnxA6 is not significantly correlated with that of Ki67 (Figure 2C-E).

**Figure 2.**
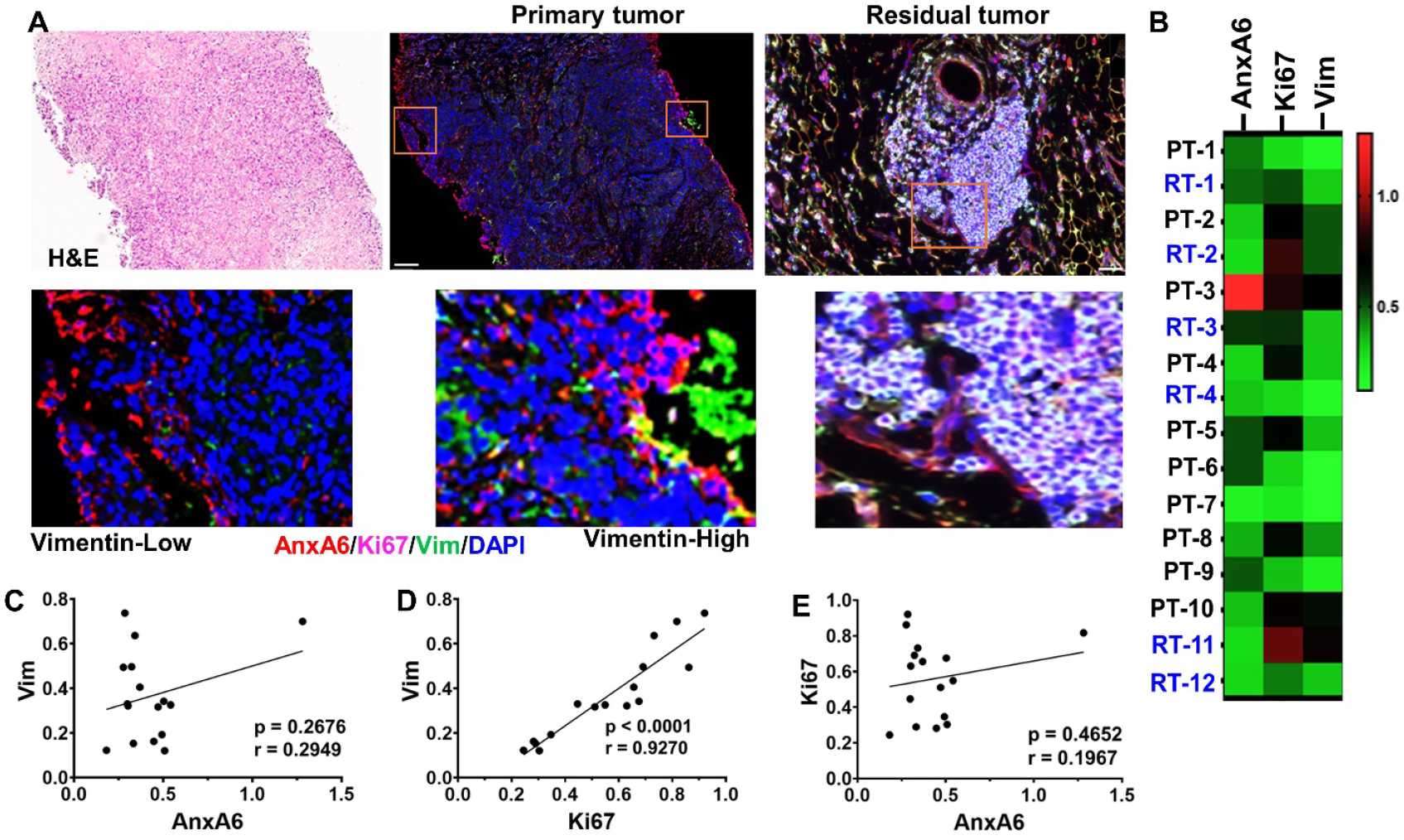
A) Expression of AnxA6 (red), Ki67 (purple) and Vimentin (green) in primary tumor versus in residual tumor tissues. A) Tissues were stained with H&E and by using tyramide based fluorescent IHC using antibodies against AnxA6 (red), Ki67 (far red), vimentin (green) and cell nuclei were stained with DAPI (blue). B) Heat map showing the relative expression of AnxA6, Ki67 and vimentin in selected biopsy (PT) and residual tumor (RT) quantified using the Ariol software (n=16). C-E) Analysis of the correlation in expression between Ki67, AnxA6 and vimentin; r denotes the correlation coefficient, p denotes statistical significance for each comparison.

### Prototype epithelial and mesenchymal-like TNBC cell lines display distinct susceptibility to standard of care chemotherapy agents

As demonstrated in previous studies (28, 30-32), the vimentin expressing BT-549 (mesenchymal-like model) express relatively high levels of AnxA6 and low levels of Ki67 (Supplementary Figure S1A and C), while the E-cadherin positive MDA-MB-468 (epithelial or basal-like model) TNBC cell express relatively low levels of AnxA6 and correspondingly, high levels of Ki67 (Supplementary Figure S1B and C). To validate this reciprocal expression of AnxA6 and Ki67, we also show that EdU incorporation in MDA-MB-468 cells was >3-fold compared to BT-549 cells (Supplementary Figure S3D-F).

We next assessed the response of the model mesenchymal-like and epithelial TNBC cells to the cytotoxic effects of doxorubicin (DOX), paclitaxel (PTX) and carboplatin (CBP). Treatment of the parental cells with various concentrations of these drugs for 72 h revealed that the epithelial MDA-MB-468 cells were 2 to 7-fold more sensitive to these drugs compared to the mesenchymal-like BT-549 cells (Supplementary Figure S1G-I). Thus, BT-549 and MDA-MB-468 cell lines are not only phenotypically distinct but are also physiologically distinct in their proliferation rates and response to standard of care chemotherapy agents.

We next established chemotherapy drug resistant populations of the mesenchymal-like BT-549 and the epithelial MDA-MB-468 TNBC cells by pulse exposure and stepwise dose escalation as described in materials and methods. Consistent with the differential response of these cells to the three chemotherapy drugs, the maximal in vitro tolerated concentrations for BT-549 cells (100 nM DOX, 30 nM PTX, and 70 µm CBP) were ∼2-fold higher than those for MDA-MB-468 cells (50 nM DOX, 15 nM PTX and 40 µM CBP). The IC^50^ values and fold resistance to these drugs revealed that BT-549 cells develop resistance to CBP and DOX but are refractory to PTX (Supplementary Figure S2A-C, Table 1), while MDA-MB-468 cells effectively developed resistance to all three chemotherapy agents (Supplementary Figure S2D-F, Table 1).

**Table 1.** Half-maximal inhibitory concentrations of standard chemotherapy drugs in parental and drug-resistant TNBC cells.

| | Half-maximal inhibitory concentrations (IC<sub>50</sub>) (μM) | | | |
| --- | --- | --- | --- | --- |
|  | BT-549-Par | BT-549-R | Drug Fold Resistance | p-value |
| **Doxorubicin** | 0.15 | 0.24 | 1.60 | 0.01* |
| **Paclitaxel** | 0.013 | 0.014 | 1.08 | 0.3391 |
| **carboplatin** | 148.5 | 336.0 | 2.26 | 0.0311* |
|  | MDA-MB-468-Par | MDA-MB-468-R | Drug Fold Resistance | p-value |
| **Doxorubicin** | 0.02 | 0.07 | 3.5 | 0.0388* |
| **Paclitaxel** | 0.06 | 0.21 | 3.5 | 0.0002* |
| **carboplatin** | 50.29 | 143.3 | 2.85 | 0.0731 |

\* Denotes statistically significant compared to parental control cells.

To confirm the differential response of these cell types to these chemotherapy agents, we compared the growth of chemotherapy drug resistant cells to untreated parental control cells cultured in ultra-low attachment plates. In agreement with the dose response curves (Supplementary Figure S2), the growth of chemotherapy resistant BT-549 cells was not different from that of parental cells (Figure 3A and B). However, for MDA-MB-468 cells, PTX-R cells grew ∼3-fold faster, while DOX-R and CBP-R cells grew slower than parental cells (Figure 3A and C). Therefore, although epithelial cells are more sensitive to these chemotherapy agents, this cell type has a higher propensity to develop resistance to any of these drugs.

**Figure 3:**
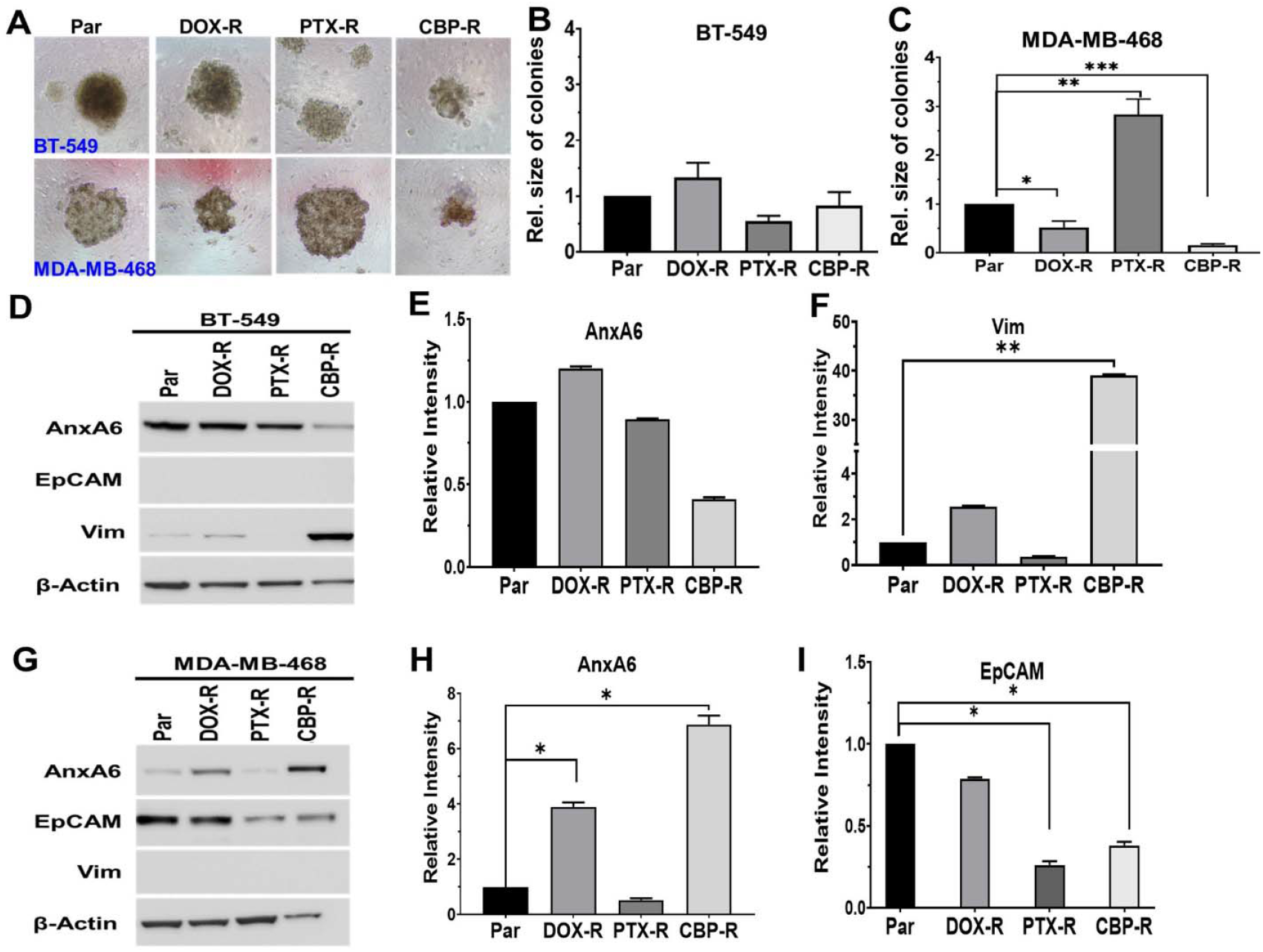
Phenotypic shift in chemotherapy resistant TNBC cells from AnxA6 and either vimentin or EpCAM in TNBC cells. A-C) Resistant mesenchymal-like BT-549 and epithelial MDA-MB-468 cells were cultured in ultra-low attachment plates for up to 10 days in medium containing the maintenance concentrations of each drug. Digital images of colonies (A) and quantification of the size of the colonies or cell masses (n>10) using ImageJ for BT-549 (B) and MDA-MB-468 (C). D and G) Cell lysates from parental and drug-resistant BT-549 (D) and MDA-MB-468 (G) cells were prepared as described in materials and methods, and the expression of AnxA6, EpCAM, and vimentin was determined by Western blotting. β-actin was used as the loading control. E, F and H, I) Densitometric analysis of protein bands using ImageJ for AnxA6 (E) and vimentin (F) in BT-549 cells and for AnxA6 (H) and EpCAM (I) in MDA-MB-468 cells relative to control cells from three independent experiments. * Denotes p<0.05; ** denotes p<0.01; *** denotes p<0.001.

### Differential expression of AnxA6 and markers of invasiveness defined chemotherapy resistant phenotypically distinct TNBC cells

We next assessed how resistance to these drugs affected the expression of AnxA6, Ki67 and vimentin, using parental untreated cells as controls. By immunofluorescence of BT-549 cells, AnxA6 was upregulated in DOX-R, unchanged in PTX-R and strongly down regulated in CBP-R cells (Supplementary Figure 3A and B). Ki67 expression decreased in both DOX-R and PTX-R but was more intense in CBP-R cells compared to parental BT-549 cells (Supplementary Figure S3A and C). Vimentin on the other hand increase in PTX-R and CBP-R cells but was unchanged in DOX-R cells (Supplementary Figure S3A and D). In MDA-MB-468 cells, AnxA6 expression was upregulated by 4-to 10-fold in the DOX-R, PTX-R and CBP-R cells (Supplementary Figure S3E and F), while the expression of Ki67 decreased in PTX-R and CBP-R but tended to increase in DOX-R cells compared to the parental cells (Supplementary Figure S3E and G). Vimentin expression also strongly increased in the resistant cells (Supplementary Figure S3E and H).

To further define the shift in the phenotype of chemotherapy resistant TNBC cells, we assessed the expression of vimentin and AnxA6 by western blotting using whole cell lysates from parental and drug-resistant cells. In CBP-R BT-549 cells, the expression of AnxA6 was downregulated while vimentin expression was upregulated (Figure 3D-F). Meanwhile, in MDA-MB-468 cells, AnxA6 expression increased in DOX-R and CBP-R cells but strongly decreased in PTX-R cells (Figure 3G and H). Expression of epithelial cell adhesion molecule (EpCAM) was strongly decreased in PTX-R and CBP-R cells and unchanged in DOX-R cells (Figure 3G and I). Consistent with heterogeneous response to various chemotherapy regimens depicted in Figure 1, PTX and CBP treatment induced opposite effects in expression of AnxA6, vimentin and Ki67 albeit in a cell type dependent manner. These data indicate that the three standard of care chemotherapy drugs induce profound phenotypic changes, particularly in epithelial cells that potentially contribute to the differential susceptibility of these cell types to chemotherapy.

### Chemotherapy resistant mesenchymal-like and epithelial TNBC cellular subsets exhibited distinct signaling patterns

To gain a better understanding of how chemotherapy resistance is sustained in epithelial and mesenchymal-like TNBC cells, we profiled the expression of 84 cancer associated proteins in cell lysates from parental control and drug-resistant BT-549 and MDA-MB-468 TNBC cells using the Proteome Profiler antibody array kit (R&D system). We identified 6 differentially expressed proteins in resistant mesenchymal-like BT-549 cells: carbonic anhydrase IX (CAIX), cathepsin D (CTSD), interleukin 6 (IL-6), CXCL8/IL-8, macrophage colony stimulating factor (M-CSF), and the inhibitor of apoptosis survivin (Figure 4A). CXCL8/IL-8 and M-CSF were the most upregulated in DOX, CBP, or PTX resistant mesenchymal-like cells. Correspondingly, the cellular levels of survivin were remarkably downregulated in PTX-R and CBP-R cells than in DOX-R cells (Figure 4A). Since most triple-negative cancers are basal-like and the majority of tumors expressing basal markers are triple-negative (33, 34), we sought to determine if these genes significantly influenced the survival of basal-like breast cancer patients using the publicly available KM Plotter tool. This analysis revealed that among the six differentially expressed proteins, low expression of survivin (Figure 4B, C), and high expression of CXCL8/IL-8 (Figure 4D, E) were significantly associated with poor relapse free survival (RFS) of basal-like breast cancer patients.

**Figure 4:**
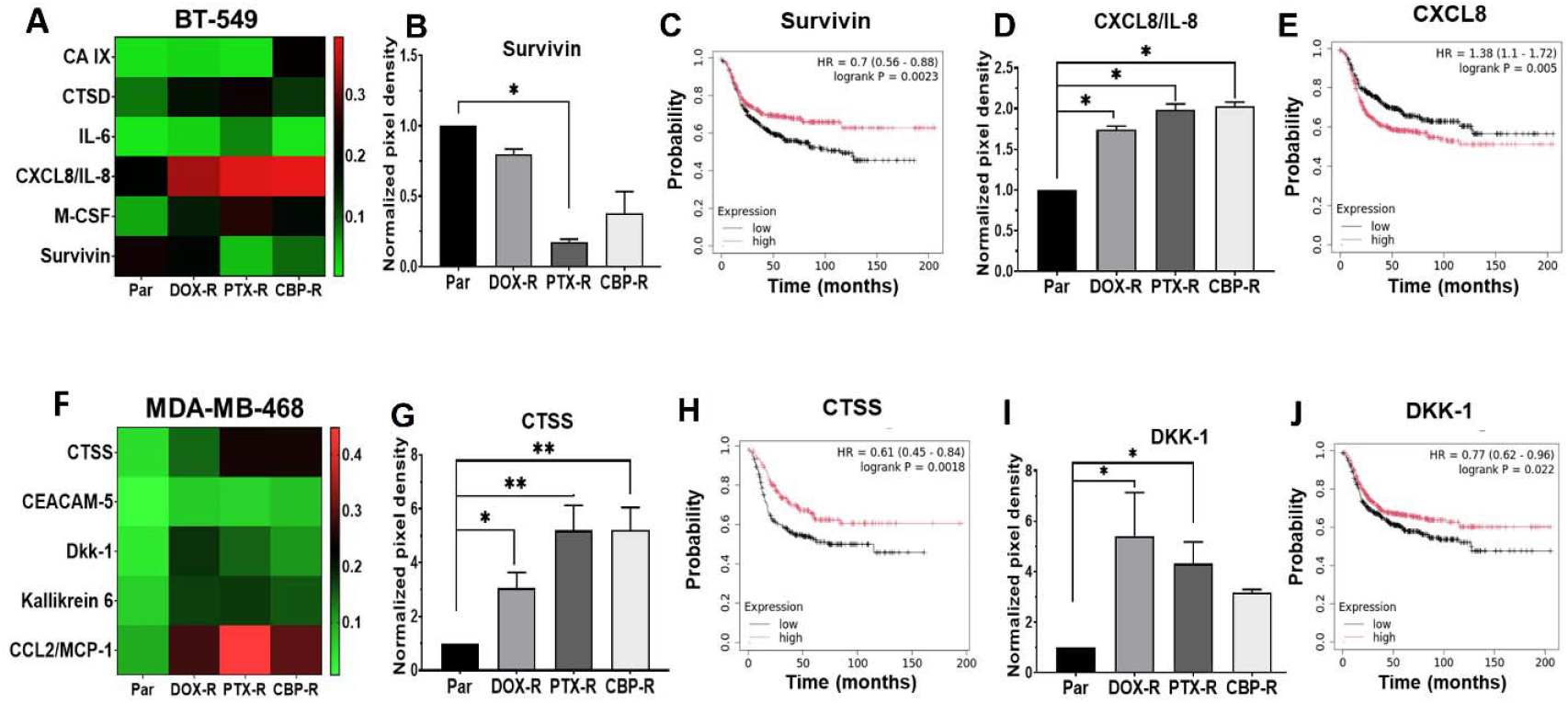
Pattern of differently expressed proteins in chemotherapeutic resistant mesenchymal-like and epithelial TNBC cells. A and F) Heatmaps showing the differentially expressed malignancy promoting proteins in cell lysates from chemotherapy resistant BT-549 (A) and MDA-MB-468 (F). Detected proteins on replica dot blots were quantified using ImageJ software and normalized to internal controls. B-E) Densitometric analysis of protein dots (B and D) from three independent experiments, and the corresponding Kaplan-Meier (KM) plots (C and E) for survivin (B and C) and interleukin 8 (E) in BT-549 cells. G-J) Densitometric analysis of protein dots (G and I) and the corresponding Kaplan-Meier (Km) plots (H and J) for CTSS (G and H) and DKK1 (I and J) in MDA-MB-468 cells. Indicated in the KM plots is the relationship between high (red) or low (black) gene expression of the indicated genes and the survival of basal-like breast cancer patients, the hazard ratio (HR) and statistical significance. * Denotes p<0.05; ** denotes p<0.01.

In the epithelial MDA-MB-468 cells, resistance to chemotherapy agents led to upregulation of monocyte chemotactic protein (CCL2/MCP-1), cathepsin S (CTSS), Dickkopf WNT signaling pathway inhibitor 1 (DKK-1), Kallikrein related peptidase 6 (KLK6) compared to parental cells (Figure 4F). As in BT-549 cells, the expression levels of these proteins were differentially influenced by the three chemotherapy agents and CCL2/MCP-1 was the most upregulated in all three drug treated cells. Patient survival analysis using the KM Plotter database revealed that the reduced expression of Cathepsin S (CTSS) (Figure 4G and H) and DKK-1 (Figure 4I and J) are significantly associated with poor RFS of basal-like breast cancer patients. Given that the identified proteins in both chemotherapy resistant mesenchymal-like and epithelial TNBC cells promote cancer metastasis via the recruitment of monocytes and macrophages into the tumor microenvironment (TME), these findings suggest that chemotherapy resistance is at least in part associated with generation of immune suppressive and tumor promoting TME by subset specific cytokines. The secretion of CXCL8/IL-8 and M-CSF by chemotherapy resistant mesenchymal-like TNBC cells as well as CCL2/MCP-1 and DKK-1 by resistant epithelial cells can signal via their cognate receptors to initiate immune cell chemotaxis and/or activate cell survival pathways.

To identify signaling pathways that potentially help maintain chemotherapy resistance in the phenotypically distinct TNBC cell subsets, we profiled 37 activated protein kinases using the phospho-kinase proteome profiler antibody array kit (R&D Systems). In the mesenchymal-like BT-549 cells, phosphorylation of Glycogen synthase kinase-3 beta (GSK-3β) at p-S9, Heat shock protein 27 (HSP27) at p-S78/S82, MSK1/2 (p-S376/360) and the non-receptor tyrosine kinase Pp60c-Src (Src) at p-Y419 were decreased, while the phosphorylation of Checkpoint Kinase 2 (CHK-2) at p-T68 increased in chemotherapy agent resistant cells compared to the control parental BT-549 cells (Figure 5A). To test whether these are clinically relevant factors in basal-like breast cancer, we show that high expression of HSP27 (Figure 5B and C) and GSK-3β (Figure 5D and E) is associated with poor survival of basal-like breast cancer patients. We also show that in the epithelial MDA-MB-468 cells, phosphorylation of GSK-3β (p-S9), p90 ribosomal S6 kinases (RSK1/2) at p-S221/S227, and Proline-rich Akt substrate 40 kDa (PRAS40) at p-T246 were remarkably increased in the PTX-R cells compared to DOX-R and CBP-R cells (Figure 5F, G and J).

**Figure 5:**
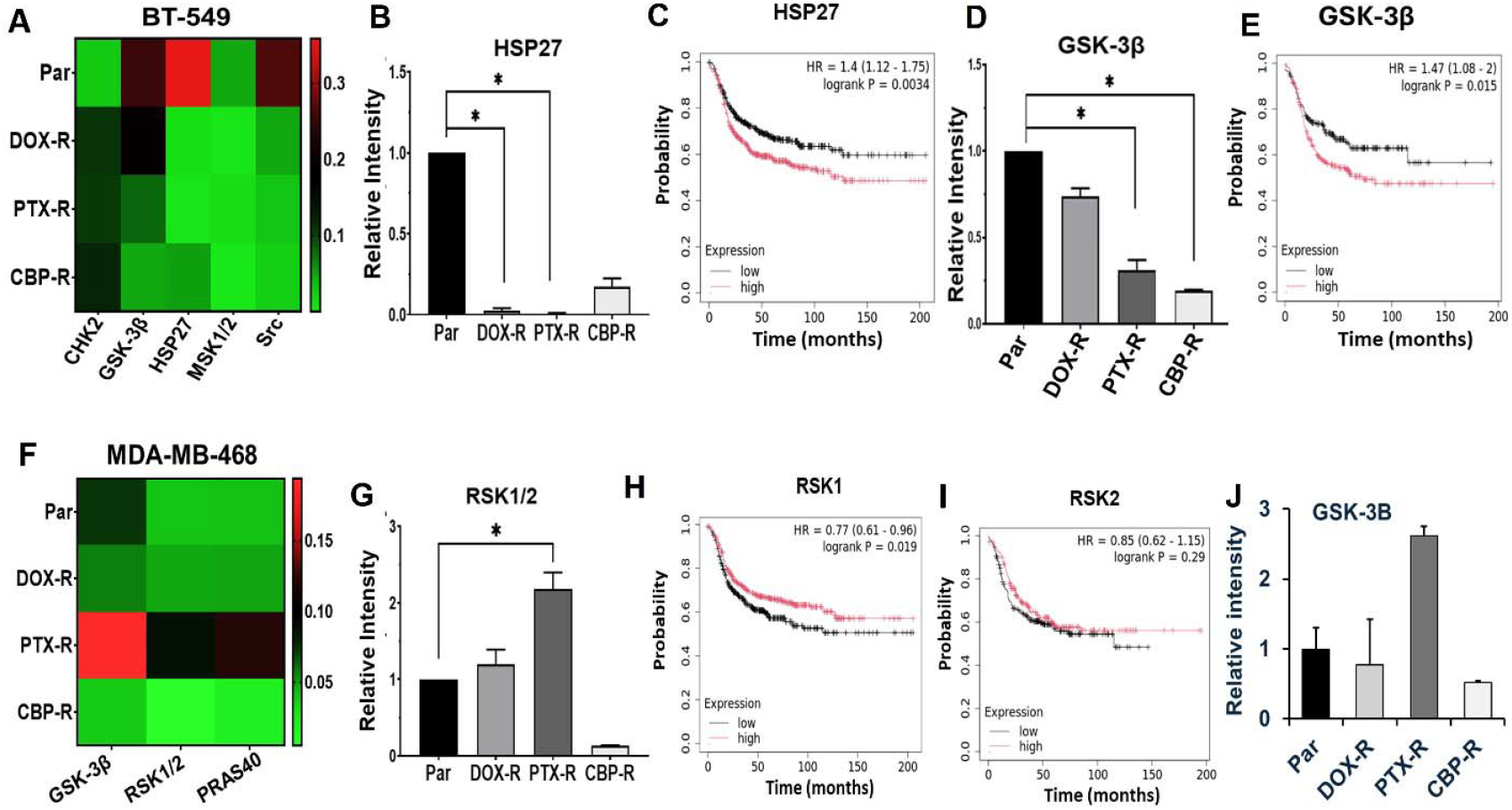
Identification of differently expressed activated protein kinases in chemotherapy resistant mesenchymal-like and epithelial TNBC cells. A and F) Heatmaps showing the differentially phosphorylated malignancy promoting kinases in cell lysates from chemotherapy resistant BT-549 (A) and MDA-MB-468 (F). Detected phospho-kinases on replica dot blots were quantified using ImageJ software and normalized to internal controls. B-E) Densitometric analysis of phosphoprotein dots (B and D) and Kaplan-Meier (KM) plots (C and E) for HSP27 (B and C) and GSK-3β (E) in BT-549 cells. G-J) Densitometric analysis of phosphoprotein dots (G and J) from three independent experiments, and Kaplan-Meier (Km) plots (H and I) for RSK1/2 (G, H and I) and GSK-3β (J and E) in MDA-MB-468 cells. Indicated in the KM plots is the relationship between high (red) or low (black) gene expression of the indicated genes and the survival of basal-like breast cancer patients, the hazard ratio (HR) and statistical significance. * Denotes p<0.05.

We next validated the expression and phosphorylation of the p90 ribosomal S6 kinase family proteins RSK1/2 and the related MSK1/2 in chemotherapy resistant TNBC cell models by western blotting. We show that RSK1/2 were expressed in both BT-549 and MDA-MB-468 cells and that phosphorylation of these proteins at S221/S227 in chemotherapy resistant cells was both drug and cell type dependent (Figure. 6A and C). Expression of MSK1/2 is restricted to the epithelial MDA-MB-468 cells and phosphorylation of these proteins at S376/360 was barely detected in DOX-R cells, and more intensely detected in CBP-R cells (Figure 6C and E).

**Figure 6:**
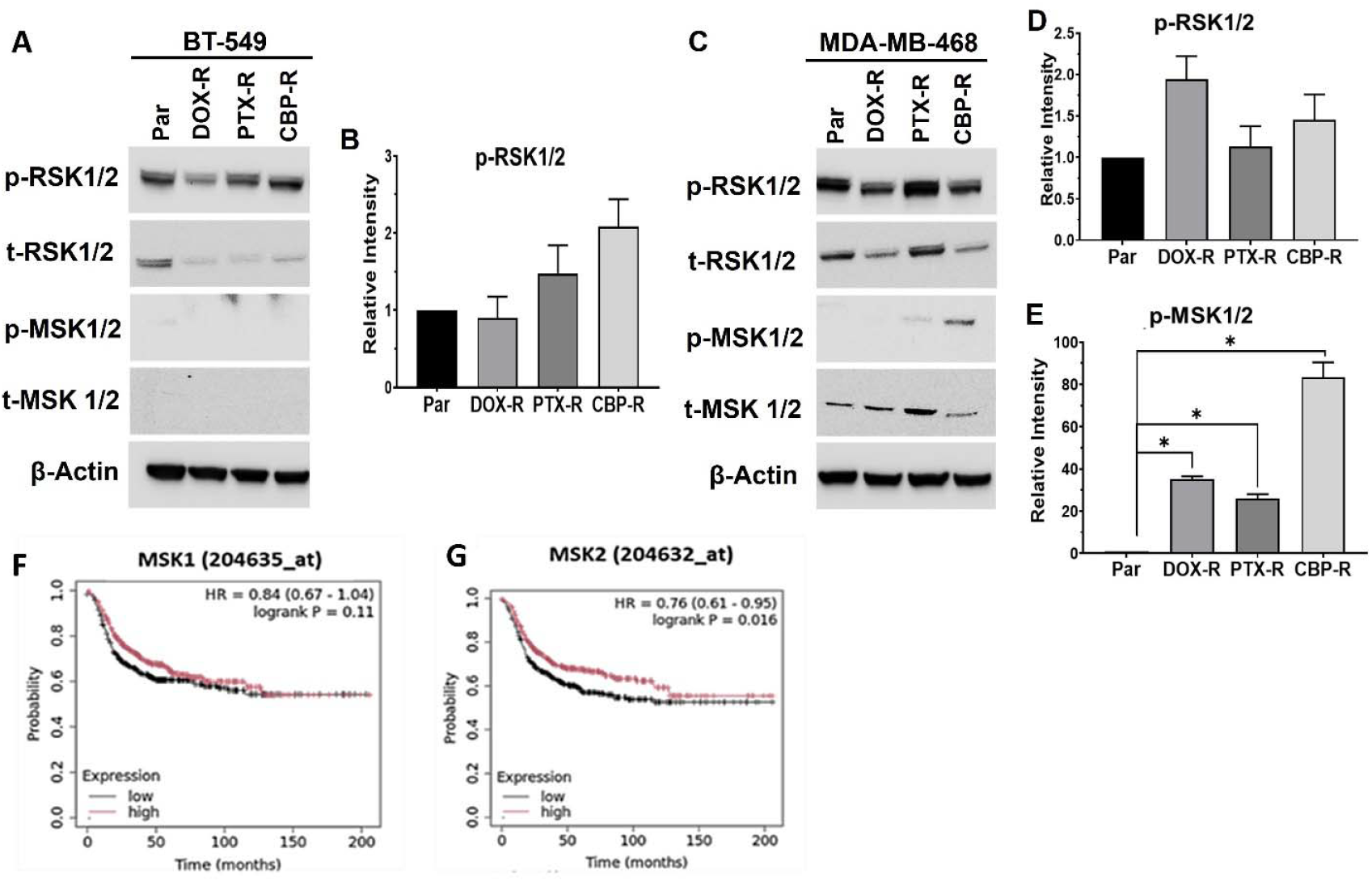
Validation of RSK1/2 and MSK1/2 phosphorylation in chemotherapy resistant BT-549 and MDA-MB-468 cells. A and C) Total and phosphorylated levels of RSK1/2 and MSK1/2 were determined by western blotting from whole cell lysates; β-actin was used as the loading control. B, D-E) Densitometric analysis of protein bands from three independent experiments for phospho-RSK1/2 in BT-549 cells (B), as well as phospho-RSK1/2 (D) and phospho-MSK1/2 (E) in MDA-MB-468 cells. F-G) Kaplan-Meier (Km) plots for RSK1 (F) and RSK2 (G). Indicated in the KM plots is the relationship between high (red) or low (black) expression levels of the indicated genes and the survival of basal-like breast cancer patients, the hazard ratio (HR) and statistical significance. * Denotes p<0.05.

Given that RSK1 and 2 as well as MSK1 and 2 proteins cannot be easily distinguished based on the molecular sizes, we again carried out KM plotter analysis of the genes to identify the clinically relevant isoforms in basal-like breast cancer. This analysis revealed that reduced expression of RSK1 but not RSK2 (Figure 5H and I), and reduced expression of MSK2 but not MSK1 (6F and G) was associated with poor RFS of basal-like breast cancer patients. Together, this indicates that a subset specific proinflammatory cytokine signature, distinct regulation of GSK-3β activity, as well as the activation of RSK1 in both cell types and MSK2 in chemotherapy resistant epithelial cells play a critical role in the maintenance of chemotherapy resistance in TNBC tumors.

## Discussion

The goal of this study was to define the contribution of mesenchymal-like and epithelial TNBC cell types in TNBC tumors to chemotherapy resistance. The most salient findings were: 1) typical TNBC tumors are predominantly epithelial tissues with discrete areas of mesenchymal-like TNBC cells, and that detection of Ki67 and AnxA6 together with EpCAM for epithelial/proliferative cells and vimentin for mesenchymal-like/invasive cells can be used to identify these cell types; 2) Resistance of TNBC cells to chemotherapy was associated with differential expression of cell type specific molecular drivers including downregulation of BIRC5 and upregulation of CXCL8/IL-8 and M-CSF in mesenchymal-like cells, and upregulation of CTSS, MCP-1 and DKK-1 in epithelial TNBC cells; 3) Differential phosphorylation of GSK-3β at S9 was increased in chemotherapy resistant epithelial cells but reduced in chemotherapy resistant mesenchymal-like cells; and 4) chemotherapy resistance in epithelial cells unlike in mesenchymal-like cells was associated with activation of MSK1/2. Overall, this study highlights the complexity and heterogeneity of TNBC tumors and reveals that the diverse response to and frequent relapse of chemotherapy residual TNBC may be attributed to the major differences in proinflammatory cytokine signaling, and activation of downstream kinases. Based on these data, we propose that chemotherapy resistant TNBCs can be alleviated by targeting these factors or the associated pathways.

The rationale to study the contribution of mesenchymal-like and epithelial TNBC cells that co-exist in TNBC tumors but with distinct drug responses is supported by previous studies that suggested that the impact of intra-tumoral heterogeneity on treatment resistance is not only defined by evolution of distinct phenotypic cell populations (16), but also how these cellular subsets with distinct intrinsic drug sensitivities co-exist within the same tumor [17]. Single-cell RNA sequencing of tumors from mouse mammary tumor virus (MMTV)-polyoma middle tumor-antigen (PyMT), MMTV-Neu, and BRCA1-null murine models of breast cancer was also used to demonstrate that these breast tumors consist of multiple breast cancer subtypes (12) that include epithelial cells and pockets of invasive mesenchymal-like cells. Also, the impact of phenotypic diversity (measured by Simpson’s score) does not always correlate with genetic intra-tumor heterogeneity (measured by the MATH index) on treatment outcomes (20). Our findings that detection of Ki67 and AnxA6 together with EpCAM for epithelial/proliferative cells and vimentin for mesenchymal-like/invasive cells can be used to identify these cell types in a typical TNBC tumor for rational treatment strategies.

Mesenchymal-like TNBC cells are inherently resistant to chemotherapy while epithelial TNBC cells are generally sensitive to chemotherapy, but residual chemotherapy resistant cells can acquire mesenchymal-like traits through epithelial-to-mesenchymal transition (EMT), that enhances their invasiveness and potential for metastasis (35). Based on the Ki67 and AnxA6 expression status, BT-549 and MDA-MB-468 TNBC cells as typical mesenchymal-like and epithelial cellular subsets respectively, indeed differed in terms of their proliferation rate, vimentin, Ki67 and AnxA6 expression and their response to chemotherapy agents. This study revealed differential cellular plasticity following prolong treatment of TNBC cells with chemotherapy agents. The drug-to-drug differences on the expression of AnxA6, Ki67, vimentin and EpCAM underscores the complexity in the treatment of TNBCs.

Our findings that vimentin expression in tumor tissues is significantly associated with Ki67 expression in TNBC patient tissues, is consistent with the previously reported association of vimentin expression with high Ki67 expression and poor prognosis for TNBC patients (36). Our data also confirms our previous report suggesting that the reciprocal expression of AnxA6 and Ki67 in TNBC tumor biopsies can delineate AnxA6-^high^/Ki67-^low^ invasive TNBC cells from AnxA6-^low^/Ki67-^high^ proliferative tumor cells (26). The observed upregulation of vimentin in carboplatin and to a lesser extent paclitaxel resistant mesenchymal-like TNBC cells and a reduction in the expression of EpCAM in carboplatin and paclitaxel treated epithelial tumor cells is consistent with invasive tendencies for TNBC cells following treatment with chemotherapy agents (37).

The development of resistance to chemotherapy has long been associated with up regulation of oncogenes and down regulation of tumor suppressor genes (38, 39), but also the modification of the tumor microenvironment to favor tumor growth and metastasis (40). This has been extensively shown to help cancer cells evade apoptosis by mechanisms that include evasion of cellular stress, enhanced DNA repair, drug efflux, and activation of survival pathways (41). Our study demonstrates that resistance of mesenchymal-like TNBC cells is associated with down regulation of BIRC5 (survivin) and upregulation of CXCL8/IL-8 and M-CSF while that of epithelial tumor cells is associated with upregulation of CTSS, MCP-1 and DKK-1. These subset specific molecular targets can block apoptosis, and signal via their cognate receptors to promote cell survival during chemotherapy. This is supported by previous studies showing upregulation of CXCL8/IL-8 in PTX resistance (42). Increased secretion of macrophage colony-stimulating factor (M-CSF) is associated with poor prognosis in various cancer types, including breast cancer and promotes tumor invasion, metastasis and chemotherapy resistance (43). Thus, upregulation of CXCL8/IL8 and M-CSF and down regulation of survivin, an inhibitor of apoptosis, clearly demonstrates the promotion of the survival of residual chemotherapy resistant mesenchymal-like TNBC cells.

The chemokine CCL2/MCP-1, has been shown to promote breast cancer progression and metastasis to lungs and bone in mouse models of breast cancer (44, 45). Cathepsin S (CTSS) has been shown to be upregulated in basal-like 1 TNBC molecular subtype with defects in DNA damage repair pathways (23) and may promote resistance to chemotherapy by facilitating the degradation of BRCA1 (46, 47). Dickkopf-1 (DKK1) is a secreted protein may promote chemotherapy resistance by inhibiting cancer cell migration and invasion via the Wnt/β-catenin signaling pathway (48, 49). Therefore, the survival and subsequent progression of chemotherapy resistant epithelial TNBC cells is at least in part promoted by DKK1, MCP-1 and the protease CTSS via inhibition of DNA repair, modulation of immune cell infiltration and proinflammatory cytokine survival pathways.

Although GSK 3β has been shown to promote chemotherapy and radiotherapy resistance as treatment options for tumors (50), it has been shown to be a tumor suppressor in some cancers including breast cancer and a tumor promoter in other types of cancers (51). The reduced inhibitory phosphorylation of GSK 3β (p-S9) in PTX-R mesenchymal-like BT-549 cells and increased GSK 3β (p-S9) in PTX-R epithelial MDA-MB-468 cells indicates significant differences in the maintenance of chemotherapy resistance in these TNBC cell types. This, at least in part, suggests stabilization of β-catenin in epithelial cells and degradation of β-catenin in mesenchymal-like cells.

Anthracyclines and platinum drugs are known to induce DNA damage which adversely affects cell cycle progression and induce apoptosis. Taxanes on the other hand affect microtubule polymerization and together can lead to serious cellular stress, disrupt protein synthesis and metabolic processes. Detection of phosphorylated mitogen- and stress-activated kinases 1/2 (MSK1/2) in MDA-MB-468 cells and phosphorylated 90 kDa ribosomal S6 kinase (RSK1/2) in both BT-549 and MDA-MB-468 cells suggest the activation of these proteins in response to metabolic and other cellular stress in chemotherapy resistant cells. Given that these proteins are very similar and can act in redundant pathways, our KM-Plotter database analysis together with western blotting confirmed that RSK1 and MSK2 are the relevant chemotherapy responsive, druggable serine/threonine RPS6KA kinase family targets in TNBC cells. Interestingly, the phosphorylation of RSK1/2 was similarly affected in the chemotherapy resistant mesenchymal-like and epithelial TNBC cells, while MSK1/2 are specifically expressed in epithelial cells and activated by prolong treatment with PTX and even more so with CBP. MSK1/2 are nuclear protein kinases that are phosphorylated and activated by MAPK/ERK pathway (52), and can promote or suppress different genes at multiple levels, which can result in the promotion or prevention of cancer metastasis (52, 53).

## Conclusions

This study highlights the complexity and heterogeneity of TNBC tumors and emphasizes the distinct characteristics and responses of mesenchymal-like and epithelial TNBC cells to chemotherapy. Chemotherapy resistance of epithelial and mesenchymal-like TNBC cellular subsets is not only associated with distinct profiles of proinflammatory and immune cell chemotactic cytokines but also modulates the activities of GSK-3β, p90 RSK1/2 and the related MSK1/2. Targeting these factors and/or the associated signaling pathways may help overcome chemotherapy resistance in TNBC.

## Supporting information

https://submit.biorxiv.org/submission/submit?roleName=author&msid=BIORXIV/2025/685128&nextpage=files&dd_disable=true

## Declarations

### Ethics approval and consent to participate

The study was conducted in accordance with the Declaration of Helsinki and approved as exempt research by the Institutional Review Board (or Ethics Committee) of Meharry Medical College (eprotocol ID 23-11-1421 and approved on 01/04/2024).

### Consent for publication

Not applicable

### Availability of data and materials

The data supporting the conclusions of this article are included within the article and its additional file.

### Author contributions

Conceptualization, NBV, AMS; methodology, NBV, MI, BRB and AMS; software, NBV; formal analysis, NVB, OKY and AMS; investigation, NVB, NIS, ARM, DB, PB, AE; writing - original draft preparation, NVB and AMS; writing – review and editing, NVB, AMS; funding acquisition, AMS and SEA. All authors have read and agreed to the final version of the manuscript.

## Acknowledgements

We thank Dr. Merry Lindsey for critically reading the manuscript, Dr. Joseph Roland, for digital histology support. We also thank the American Cancer Society for a postdoctoral fellowship to NVB; and the Meharry RCMI Research Capacity Core (MRRCC) facility for technical support. The schematic graphical abstract was created using BioRender.

