## Supplementary material for "Contribution of Mesenchymal-like and Epithelial Cellular Subsets to Chemotherapy Resistance in Triple-Negative Breast Cancer": https://submit.biorxiv.org/submission/submit?roleName=author&msid=BIORXIV/2025/685128&nextpage=files&dd_disable=true

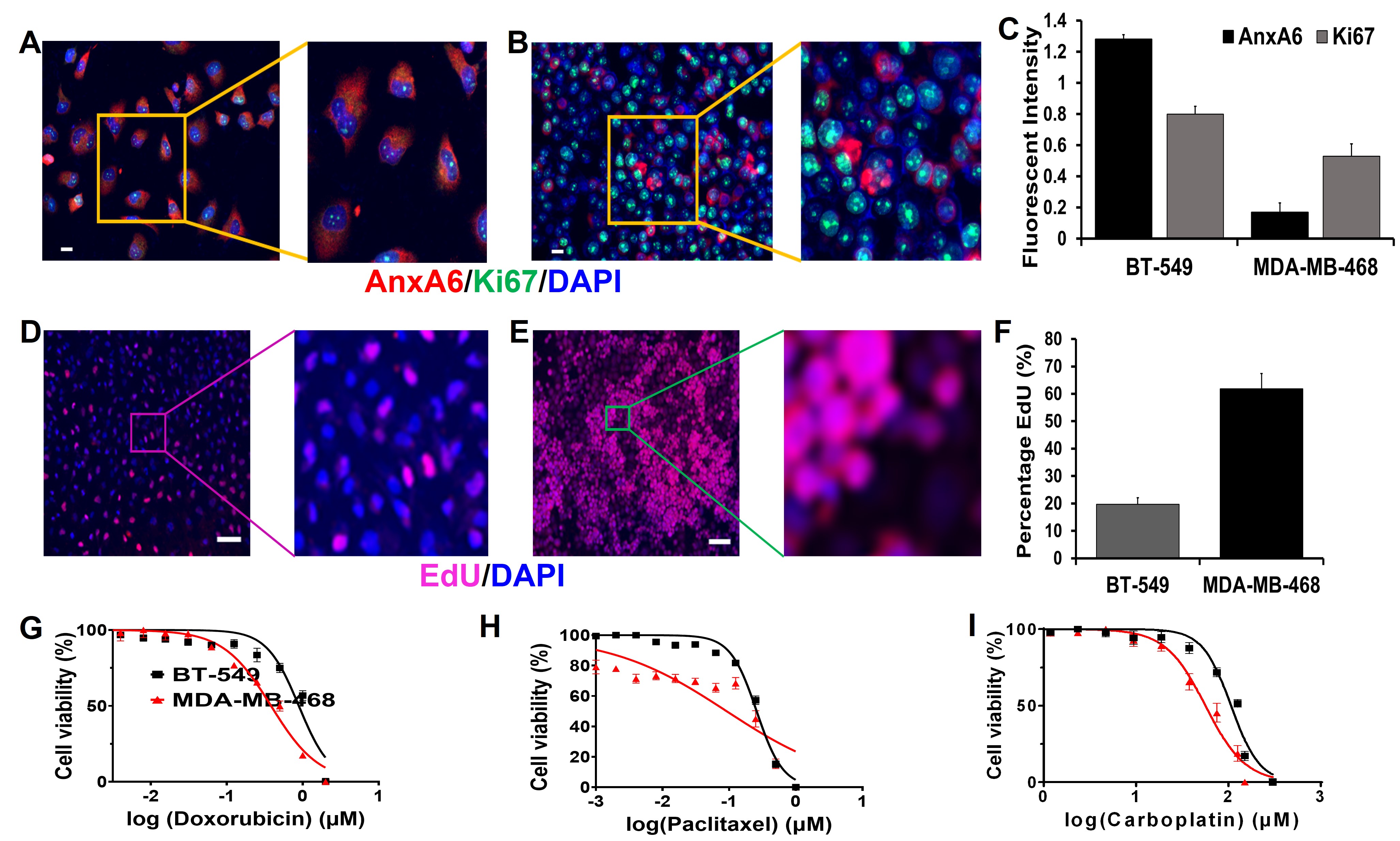

**Supplementary Figure S1. Validation of model in epithelial and mesenchymal-like TNBC cells.** A-B) by immunocytochemical staining of AnxA6 (red), and KI67 (green) and cell nuclei with DAPI (blue) in the mesenchymal-like BT-549 (A) and the epithelial MDA-MB-468 (B) TNBC cells. C) Quantification of the fluorescent intensity of AnxA6 and Ki67 in the TNBC cell lines. Data are expressed as mean ± SD (n = 3/group). D-F) Analysis of cell proliferation by DNA synthesis using the Click-EdU Assay. Images represent cells labeled with EdU and stained with anti-EdU (red), and cell nuclei with DAPI (blue) in the mesenchymal-like BT-549 (D) and the epithelial MDA-MB-468 (E) TNBC cells. F) Quantification of EdU staining. Data are expressed as mean ± SD (n = 3/ group). G-I) Response of the mesenchymal-like BT-549 and the epithelial MDA-MB-468 TNBC cells to doxorubicin (G), paclitaxel (H) and carboplatin (I). Cell viability was assayed by using the PrestoBlue reagent. Data were collected as quadruplicates each time and experiments were performed twice.

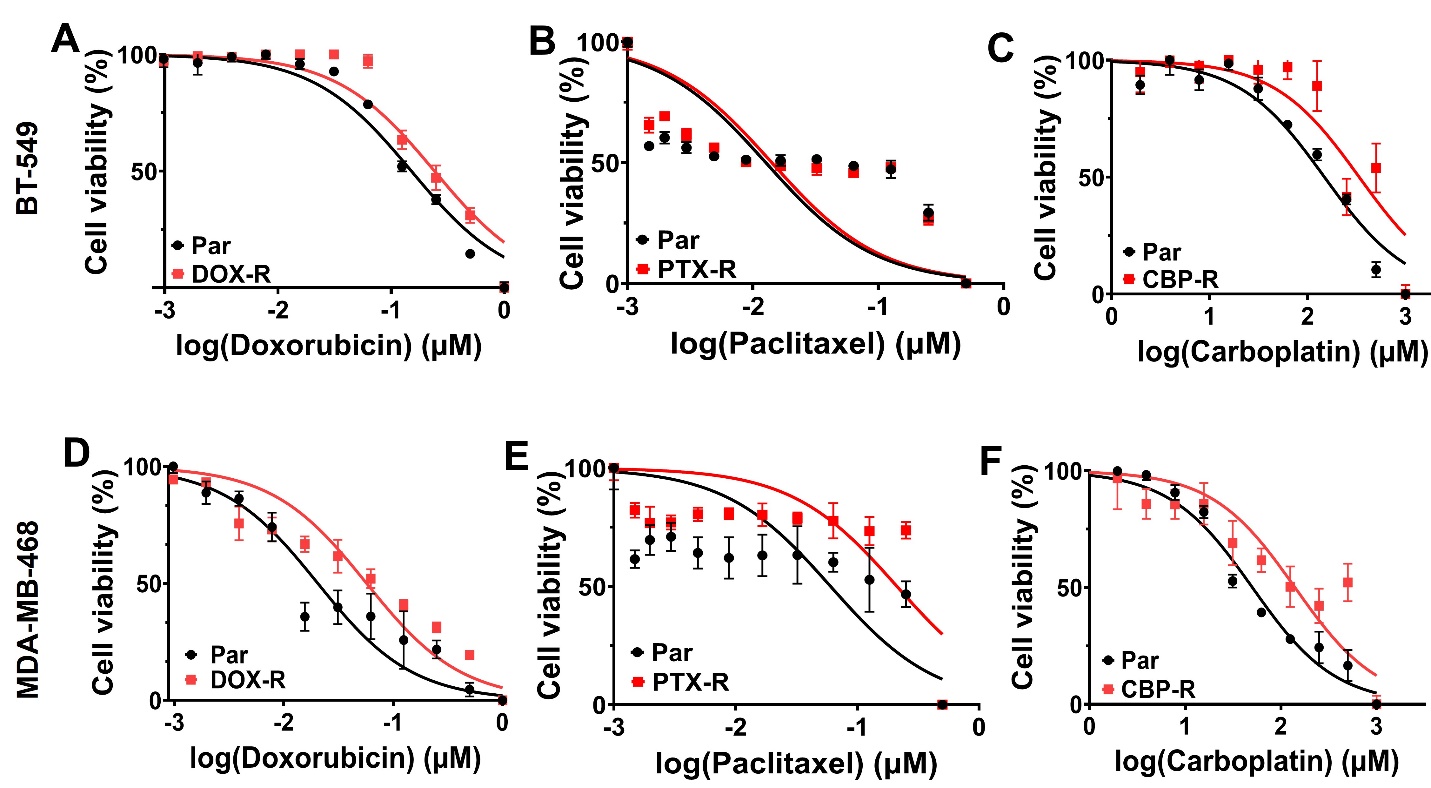

**Supplementary Figure S2. Chemotherapy resistance induced phenotypic changes in mesenchymal-like and epithelial TNBC cells.** A-F) Dose-response curves of resistant BT-549 cells (A-C) and MDA-MB-468 cells (D-F) to standard of care chemotherapy agents. Cell viability was assayed by using PrestoBlue reagent. Data were collected as quadruplicates and experiments were performed twice.

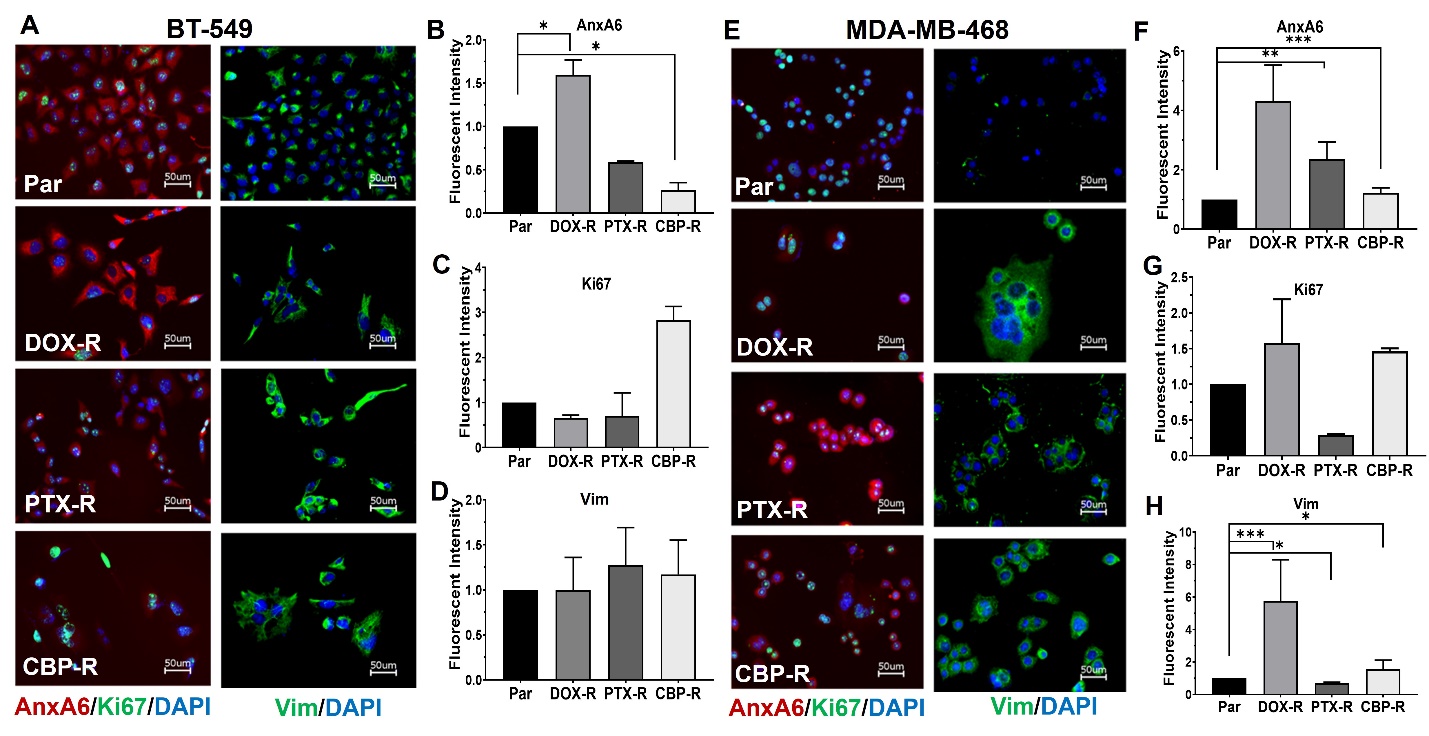

**Supplementary Figure S3: AnxA6, Ki67 and vimentin in chemotherapy drug resistant TNBC cells.** A and E) Representative images of the expression of AnxA6 (red), Ki67 (green) and vimentin (green) in control (Par) and chemotherapy drug resistant BT-549 (A) and MDA-MB-468 TNBC cells (E), cell nuclei were stained with DAPI (blue). B-D) Quantification of the fluorescence intensity of AnxA6 (B), Ki67 (C), and vimentin (D) in parental and dug resistant BT-549 cells; and of AnxA6 (E), Ki67 (F), and vimentin (G) in parental and dug resistant MDA-MB-468 cells. Data are expressed as mean ± SD (n = 3/group). * Denotes p<0.05; ** denotes p<0.01; *** denotes p<0.001.
